# Estimating global enzyme abundance levels from cofactor requirements: a model-based analysis of the iron metabolism in yeast

**DOI:** 10.1101/229104

**Authors:** Duygu Dikicioglu, Stephen G. Oliver

## Abstract

Metabolic networks adapt to changes in their environment by modulating the activity of their enzymes and transporters; often by changing their abundance. Understanding such quantitative changes can shed light onto how metabolic adaptation works, or how it can fail and lead to a metabolically dysfunctional state. Unfortunately such data are only available on a fraction of the enzyme and transporter pools, and this leaves us distant from generating the full picture. We here propose a strategy to quantify metabolic protein requirements for cofactor-bearing enzymes and transporters through constraint-based modelling. In this work, we constructed the first eukaryotic genome-scale metabolic model with a comprehensive iron metabolism and investigated partial functional impairment of the genes involved in iron-sulphur (Fe-S) cluster maturation, which revealed extensive rewiring of the fluxes in silico, despite only marginal phenotypic dissimilarities. The yeast metabolism at steady state was determined to employ a constant iron-recruiting enzyme turnover at a rate of 3.02 × 10^−11^ mM / (g biomass)^−1^ h^−1^. Furthermore, we employed this model to identify and study a mutualistic relationship between *Streptomyces coelicolor* and *Saccharomyces cerevisiae*, which involves iron and folate sharing economies between the two microbes.

## Introduction

The complex network structure comprised of the interactions among enzymes, metabolites and their regulators define the phenotypes of all biological systems [1]. Two sets of components of this network structure, the enzymes and the metabolites comprise the metabolic network of an organism. The fine-tuning between enzymes and metabolites has been quantitatively shown to modify metabolic reaction rates; enzyme abundance and metabolite concentration was identified to inversely act upon limiting metabolic reaction rates in *Saccharomyces* (*S.*) *cerevisiae* [2], Consequently, understanding these relationships can help us address the nature of the transition between metabolic states. Metabolic state shifts frequently demand improved adaptive response of the organism to environmental or genetic perturbations. While this response can be a desirable trait in many biotechnological applications, it needs to be overcome in some instances, for example, in drug resistance development.

The measurement of intracellular metabolite pools and protein abundances, as well as understanding the relationships between the two has been of interest for many different systems ranging from *Escherichia coli* [3] to tumour cell lines [4,5], due to their pivotal role in understanding and tuning metabolic adjustment. Total protein content and absolute quantification of individual proteins were reported for a number of biological systems [3-7] to assist these efforts. However, the available methods cannot focus on studying enzymatic proteins in isolation. Enzymes are generally present in lower copy numbers than many other types of proteins constituting the proteomic landscape, such as ribosomal proteins. Furthermore, some enzymes are present only in very low copy numbers pushing the limits of detection and quantification of available analytical techniques [6]. A recent study on the direct and absolute quantification of over 1800 yeast proteins revealed that only less than a quarter of the quantified proteins were indeed metabolic [8].

Metabolic models with high predictive ability have the potential to serve as ideal workhorses for investigation of the metabolic network and its components [9]. Metabolic models can provide reasonable predictions when actual measurement of network components is not feasible, or even possible, as demonstrated by predictions on flux distributions at genome-scale [10]. Quantitative predictions on absolute fluxes may serve as proxy for the relative abundance of the enzymes catalysing those reactions with the predicted flux. However, this is still far from providing an absolute measure, which can then directly relate to the substrate and enzyme requirements of the involved reactions.

We propose here a network-based strategy to determine optimal enzyme abundance by model predictions based on the metabolic requirements of their cognate cofactors. We study a family of non-metabolic intermediate cofactors that do not have donor functional groups. The iron cofactor family including iron in its ionic form (Fe^3+^, Fe^2+^) and complex ion entities such as the haem family (sirohaem, haems A, B, C and 0), and the iron-sulphur clusters (2Fe-2S, 4Fe-4S) were the choice of interest in this study as these iron-containing cofactors bind nearly 10% of the documented enzymes and transporters in the metabolism. Baker’s yeast, being the first eukaryote with a complete genome sequenced [11] has been a favourite model organism for investigating molecular mechanisms in higher eukaryotes including humans [12], Cellular activities including DNA replication, recombination and repair, RNA transcription and translation, intracellular trafficking, as well as the enzymatic activities of the general metabolism, and mitochondrial biogenesis are conserved from yeast to human [13]. The availability of comprehensive and powerful genome-scale models of the yeast metabolism [9] for nearly the past two decades [14] rendered yeast model ideal for this study. Although there is an extensive wealth of knowledge regarding the iron utilization and homeostasis in yeast [15-17], the integration of this information with that of the curated genome-scale metabolic network has not yet been achieved, limiting its practice in in silico studies. This issue has recently been brought to attention in a comprehensive study on the comparative analysis of yeast metabolic models [18].

In this work, we extended the genome-scale metabolic network model of yeast to incorporate iron metabolism, and made the first comprehensive mathematical representation of any inorganic ion entity in a eukaryotic model organism network available for the community (BioModels [19] MODEL1709260000). We have benchmarked the new model against environmental and genetic modifications, and have verified cellular responses against availability of iron and copper, and also identified extensive re-wiring of the fluxes to cope with the lowering of the dosage of the essential Fe-S cluster maturation genes. The model allowed us to establish the iron family cofactor requirements based on how the fluxes were distributed through the metabolic network. This requirement was then employed as proxy for calculating the metabolic requirement of those enzymes and transporters of the metabolism, which employ iron species as cofactors. We finally report a potential commensal symbiosis between *S. cerevisiae* and *Streptomyces (Str.) coelicolor* suggested by this network-based analysis of the metabolism, which involved the transaction of folate from the yeast to the bacterium in exchange for the provision of siderophore-bound iron by the bacteria to be at yeast’s disposal.

## Materials and Methods

### Modelling Methods

#### Primary metabolic model, simulation environment, and model annotation

The primary model for the incorporation of the iron metabolism was selected as the most recent stoichiometric model of the S. cerevisiae metabolic network (v7.6) [9]. The extended model (Yeast7.Fe) is provided as Supplementary Material 1 and the details on modification and extension of the existing model are provided in Supplementary Material 2. The extension annotations were documented in compliance with the MIRIAM standards [20]; ChEBI [21] and KEGG [22] identifiers (IDs) of the compounds, KEGG [22] and SGD [23] IDs of the genes, UniProt [24] IDs of the enzymes, and KEGG [22] IDs of the reactions were documented whenever applicable. PUBMED IDs [25] of the relevant publications were provided as the resource if the KEGG reaction IDs were not available. Logic rules for multiple genes involved in the catalysis of a single reaction were provided as documented in their cognate literature (Supplementary Material 2).

The extended genome-scale model of S. cerevisiae is available in a COBRA compatible-SBML format (v.4). Model simulations were carried out in MATLAB environment R2016b (9.1.0.441655, Mathworks, USA) with the COBRA Toolbox (v3.1.2), the SBML Toolbox v4.1.0 and libSBML library V5.15.0 running in the background employing standard linear optimization techniques (Gurobi 7.0.2). The objective function was to maximise growth. A specific growth rate of 0.1 h^−1^ was used unless otherwise specified.

#### Cofactor representation

For any metabolic reaction catalysed by a cofactor requiring enzyme (N):

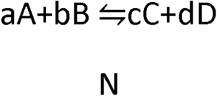

where the stoichiometric coefficients of the substrates (A and B) and products (C and D) are denoted by their cognate lower case letters, a cofactor (X) of enzyme N enters the metabolic reaction as a substrate and leaves as a catalytically unreactive product:

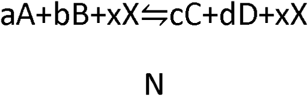

Iron ions, copper ions, pyridoxine, haem entities and Fe-S clusters were integrated as cofactors (X) in the model. The relative amount of cofactors required for each reaction was designated by x. The number of active sites on N was assumed to be 1 in all cases due to the unavailability of data, and this factor of overestimation was knowingly introduced into the model.

#### Evaluation of the predictive power of the model

Prediction of gene essentiality was used as the evaluation criterion for the new reconstruction. Empirical data on S. cerevisiae S288C strain essentiality was obtained from Saccharomyces Genome Database (SGD) [23] (website accessed on 22/03/2017). *IMA3* and *IMA4* were excluded from analysis as these genes hold 100% similarity. A full list of non-SGD resources for essentiality is provided in Supplementary Material 3. A Boolean criterion was adopted for the presence or absence of growth, where a positive or a negative call were defined as the presence (= 1) or absence (= 0) of growth, respectively. A true positive (TP) was a non-essential gene deletion mutant predicted as viable and a true negative (TN) was an essential gene deletion mutant predicted as inviable. A false positive (FP) designated a viable prediction for the deletion of an essential gene, whereas a false negative (FN) was nominated when the model yielded an inviable prediction for the deletion of a nonessential gene. The following success measures were employed to assess the predictions: positive predictive value (PPN)= TP/(TP + FP), negative predictive value (NPN)= TN/(TN + FN), sensitivity= TP/(TP + FN), specificity= TN/(TN + FP), and predictive success= (TP + TN)/(TP + TN + FP+ FN). Hypergeometric p-values were determined based on the cumulative distribution functions for determining under- or over-enrichment of factors.

### Experimental Methods

#### Cell line, cultivation conditions, subcellular fraction enrichment protocols

Heterozygous deletion mutants *HO/Δho*, *ARH1/Δarh1*, *ARH1/Δarh1* and *YFH1/Δyfh1* of *S. cerevisiae* BY4743 (background: *MATa/α his3Δ1/his3Δ1 leu2Δ0/leu2Δ0 lys2Δ0/LYS2 MET15/met15Δ0 ura3Δ0/ura3Δ0*) strain were employed in this study [26]. Deletion of a single copy was verified by PCR using the confirmation primers [26]. Qiagen DNeasy Blood & Tissue Kit was used for isolation and purification of DNA from the cell extracts as described in the Kit Protocol.

Three separate cultures were grown at 30°C in YPD media allowing sufficient aeration in vented cap low protein binding tissue culture flasks (TPP^®^) (surface area (cm^2^) to height ratio (cm) = 300:4.5) with shaking (220 rpm) to an OD_600_ of 0.7. Fresh medium from the same batch was employed in further analytical assays. Supernatant was collected by centrifugation at 7.6k rpm for 10 min at 4°C. The cell wall digestion and lysis of the harvested cells (1 OD_600_ equivalent) was carried out as described in Qiagen DNeasy Blood & Tissue Kit protocol, and crude cell extracts as well as the supernatant was stored at −20°C. Yeast Mitochondria Isolation Kit from Sigma-Aldrich (Cat no: MIT0IS03) was used for the isolation of an enriched mitochondrial fraction from yeast cells starting from 20 OD_600_ equivalent culture and intact mitochondria was stored −20°C until further use. Cytosol was separated from the particulate organelles in the cell through an oil layer employing abcam’s Cytosol/Particulate Rapid Separation Kit (Cat no: ab65398) from a 0.6 OD_600_ equivalent culture. A vacuole-enriched fraction was isolated from 420 OD600 equivalent yeast cells following a yeast-specific protocol as described [27], Proteins in all fractions were precipitated in 20%w/v trichloroacetic acid at 4°C [28], and protein free lysates were analysed separately.

#### Analytical Assays

The glucose, glycerol, ethanol and ammonium content of the supernatant were determined enzymatically by r-biopharm Roche Yellow Line assays (Cat no: 10716251035, 10148270035, 10176290035, and 11112732035, respectively). Copper content, the total haem content, and the Fe2+ and total iron content of the samples post reduction were determined colorimetrically employing the Copper Assay Kit by Sigma-Aldrich (Cat no: MAK127), BioAssay Systems QuantiChromTM Heme Assay Kit (Cat no: DIHM-250), and the Iron Assay Kit by Sigma-Aldrich (Cat no: MAK025), respectively, as described by their manufacturers. Assays were executed in 96-well Corning^®^ Costar^®^ 96-Well flat–bottom Cell Culture Plates whenever applicable. All data pertaining these analyses are provided in Supplementary Material 3.

## Modelling and Implementation

### Iron metabolism in yeast metabolic models

Although the predictive power of yeast metabolic network models improved substantially, the predictive capability of these models remains limited by missing and incomplete representation of metabolic pathways in the model [29]. Our previous work on Yeast 7 metabolic network model highlighted the high metabolic burden carried by the energy generation linked pathways and processes imposing limitations on the model’s predictive ability [29]. A recent analysis we carried out by constraining this model by the intracellular fluxes of the purine nucleotide biosynthetic pathway intermediates determined by HPLC analysis [30] demonstrated that the prediction of growth rate was at least 100% off from the experimentally determined value. The distribution of the fluxes indicated that iron uptake and utilization pathways were inactive as they were inadequately represented and completely disconnected from the rest of the metabolic network (Figure 1a). We addressed all such problems through extensive literature curation and explain how the iron metabolism was reconstructed into a network in the following sections.

**Figure 1.**
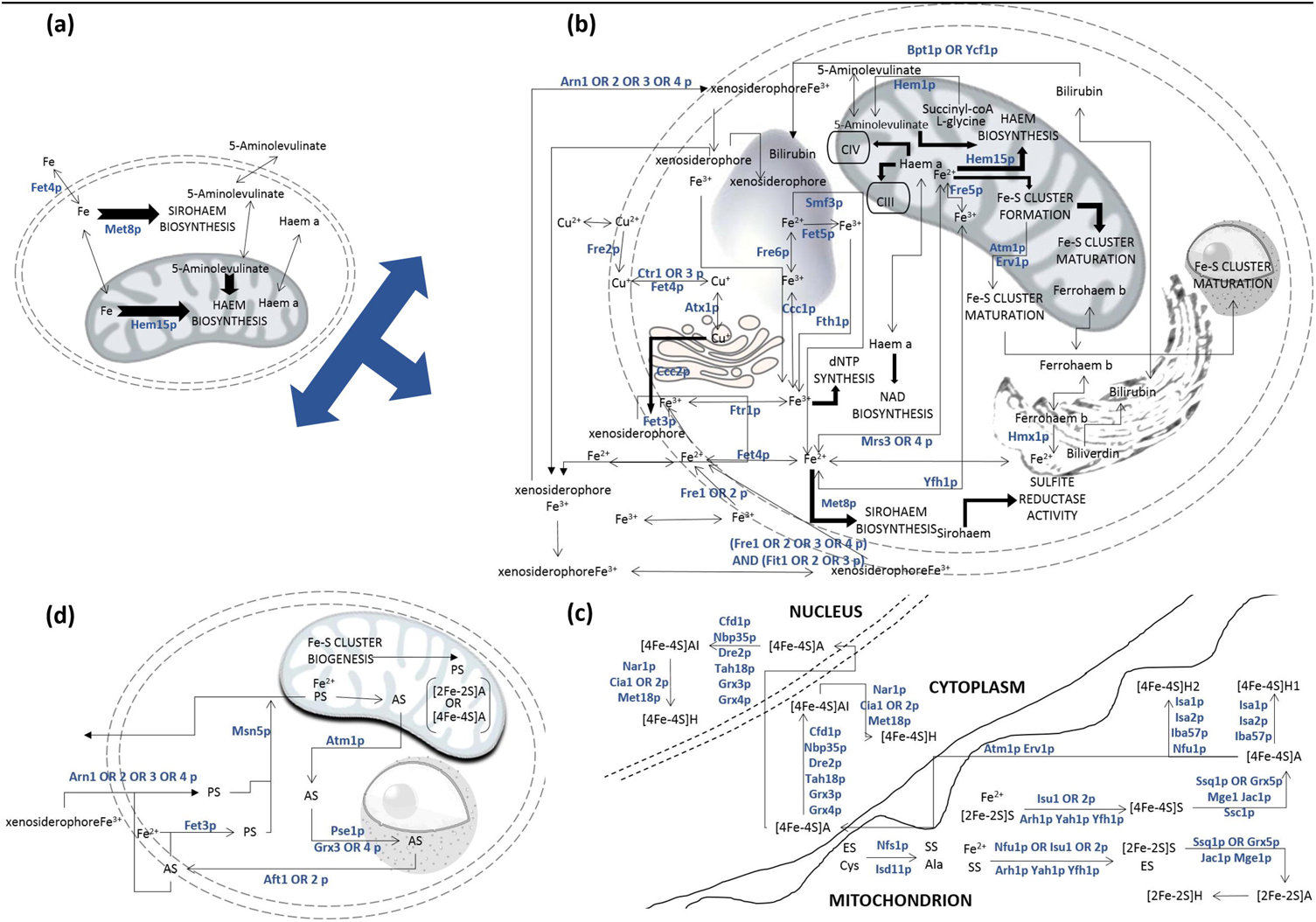
Schematic representation of iron metabolism in the yeast model. Pathways and metabolites are designated in uppercase and lowercase letters, respectively. Directionality of the fluxes through the pathway is specified by the arrows (for single reaction steps) and block arrows (for lumped consecutive reaction steps). Metabolic enzymes are given in teal colour. The cell boundary and organelle boundaries are represented as double dashed; the mitochondrion, nucleus, vacuole and ER have cartoon representations. The primitive representation of iron metabolism in the existing Yeast 7.6 model is provided in (a). The details on the reductive, non-reductive, and xenosiderophore-bound iron uptake, intracellular transport and storage of iron, haem and sirohaem biosynthetic and degradation pathways are provided in (b). Details regarding the biogenesis of Fe-S clusters in the mitochondrion (ISC machinery), and the maturation of apoenzymes (A) into Fe-S cluster-bound holoenzymes (H) in the mitochondrion (ISC machinery), in the cytosol, and in the nucleus (CIA machinery) are provided in (c). An empty scaffold (ES) and its sulfonylated form (SS) were introduced as pseudo-metabolites in Fe-S cluster formation mechanism. The regulation of iron uptake via the iron regulon employing the negative feedback from Fe-S cluster biogenesis is demonstrated in (d). The shuttling of the signals representing the availability of mitochondrial iron (PS) and its depletion (AS) were introduced as pseudo-metabolites to modulate and activate the reductive iron uptake routes. For simplifications on the function and activity of the iron regulon, see Text.

### Design considerations – 1: Uptake and intracellular transport and storage of iron

Both reductive (Fe^3+^ - transporting, high affinity) and non-reductive (Fe^2+^ - transporting, low affinity) iron uptake mechanisms were incorporated in the metabolic network. Free and xenosiderophore- bound high-affinity iron uptake was represented by the reductive pathway. The intracellular transport of iron and complex iron entities such as haem and sirohaem across intracellular boundaries and the storage of residual components and excess iron were also considered Figure 1b). Iron export was not evidenced in *S. cerevisiae* [31], and was consequently excluded.

The lower bound of the flux through the low-affinity iron uptake reactions was set at 1μM [32], The high and low affinity uptake systems were modelled to work exclusive of each other, if needed [31,32], Specific ARN family transporters involved in the uptake of each iron-bound xenosiderophore and the fate of each xenosiderophore in the yeast cell was modelled individually for ferrichrome, N,N',N"-triacetylfusarinine C (TAFC), enterobactin and ferrioxamine B (Supplementary Material 2) [31]. The reductive assimilation of iron bound to xenosiderophores was facilitated by one of the functionally non-interchangeable metalloreductase (Frelp – 4p) [33] and all of a family of mannoproteins (Fit1p – 3p) that are incorporated into the cell wall via glycosylphosphatidylinositol anchors in *S. cerevisiae* [31]. The copper recruitment of the reductive pathway of high-affinity iron uptake [34] necessitated the incorporation of copper uptake in the metabolic network. Since copper metabolism was not represented in the primary metabolic network of yeast to any degree, the network was extended to incorporate the uptake of copper and its function in high affinity iron uptake. It is important to note that the representation of copper was not exhaustive; only activities that are relevant for the iron metabolism were considered in this reconstruction. The threshold for switching between the high and low affinity copper transport was set as 20μM [35]. Two assumptions were made in modelling the uptake of copper across the cell envelope: (i) Although included as a unique species in the model, the high affinity copper transporter Ctr3p was not associated with the copper uptake reaction in the model along with Ctr1p, since the Ty2 transposon in S288C strains employed in this study rendered the protein inactive [36]. (ii) Fre1p copper reductase was similarly excluded from gene-reaction associations since it was reported to be active only during the first 3-4 hours of culture growth [37], The *in silico* analyses conducted in this study are all carried out under pseudo- steady state assumption, and by that time Fre2p was thought to govern copper reductase activity.

### Design considerations – 2: Biosynthesis and recycling of complex iron entities

The biosynthesis and recycling of haem, sirohaem and Fe/S clusters were fully implemented in the genome scale model of the yeast metabolic network. 5-aminolevulinate synthesis from glycine and succinyl-CoA via the Shemin pathway [38] was introduced and 5-aminolevulinate uptake was excluded from the model. Haem catabolism in the endoplasmic reticulum, and the storage of bilirubin in the vacuole were also introduced to the model *de novo* (Figure 1b).

A pseudo-metabolite; mitochondrial empty scaffold (ES) was introduced to the metabolic network in order to model the Fe-S cluster biogenesis (the ISC machinery) in the mitochondria and the Fe-S cluster assembly (the CIA machinery) in the cytosol and the nucleus, which allowed the maturation of the Fe-S clusters and their incorporation on their cognate apoenzymes. Although this scaffold itself was thought to be a transient protein complex [16], the members of this complex were associated with the “reaction” step in the metabolic model converting an empty scaffold into a sulfonylated scaffold (SS) to render the process similar to a reaction. The ES was then released in the next step once the Fe-S cluster itself was formed. 2Fe-2S and 4Fe-4S cluster maturation was assigned to the mitochondrion, the cytosol or the nucleus as applicable. Two different mitochondrial 4Fe-4S maturation routes were implemented to account for the genetic aspects of 4Fe-4S cluster maturation for Lip5p and Sdh2p or for other mitochondrial Fe-S proteins (denoted as H1 or H2 in Figure 1c, respectively) [16].

### Design considerations – 3: Iron regulon

Two different regulatory mechanisms for iron uptake via the iron regulon were implemented in this model: (i) the yeast-specific haem-regulated positive feedback route [16], and (ii) the mitochondrial Fe-S cluster biogenesis-associated negative feedback route.

Haem deficiency, indicated by the deactivation of the enzymes encoded by the essential genes (*HEM1-4*, *HEM12*, *HEM13*, *HEM15*) of the pathway, was reported to be an indicator of low iron uptake in yeast [16]. We coupled the essential enzymes of haem biosynthesis to the reaction representing low affinity iron uptake catalysed by Fet4p to account for this positive feedback mechanism. This coupling with low-affinity iron uptake ensured that haem biosynthesis would not be over-activated by the network unless extracellular iron was abundant.

The negative feedback on iron uptake by Fe/S cluster biogenesis being shut down upon iron depletion was modelled by introducing two new pseudo-metabolites: AS signalling the depletion of intracellular iron, and PS indicating the intracellular availability of iron (SBO:0000409 term: interaction outcome). AS activated the iron regulon in the nucleus, relaying a message to the cell envelope to activate the uptake of iron (Figure 1d). PS was coupled with the formation of Fe-S apoclusters in the mitochondrion. Unavailability of iron produced AS to be then relayed from the mitochondrion to the nucleus and further to the cell envelope, unchanged. Msn5p, a karyopherin shuttling between the nucleus and the cytoplasm, was assigned a modified function in the model, being associated to the transport of PS from the cell envelope back to mitochondrion designating the relay of the signal for the presence of iron in the cell. Any signal relaying event was designated by the SBO:0000464 term: “state variable” in the model. Aft1p, which was responsible for relaying the iron depletion signal to the cell boundary, was reported to have an additional role in creating iron resources for the cell by binding Cth2p to facilitate the mRNA degradation of iron-containing enzymes [39]. This route was excluded from the network as enzymes were never treated as substrates or products in the current model. Aft1p was also reported to mediate saving iron by activating Vth1p-mediated biotin uptake since biotin synthesis was reported to be iron consuming [40]. This route was also excluded as biotin co-enzyme metabolism was not considered directly as our system of interest.

### Iron family cofactor considerations of metabolic enzymes

Iron family cofactors, copper ions and pyridoxine; in one affiliated reaction, were employed as a substrate and untransformed reactant in those reactions that were catalysed by enzymes activated by these cofactors, as described in ‘Cofactor representation’. This allowed to establish the connectivity between the iron metabolism, described above, and the primary metabolic network of yeast. For this purpose, the copper, iron, haem, sirohaem, pyridoxine and Fe-S requirements of metabolic were identified (Supplementary Materials 4, 5) [17,41-43]. This compilation excluded 41 enzymes similarly reported to have iron cofactor involvements due to several reasons; the documented information was observed to conflict with other available information; the enzyme was not included in the metabolic network model Y7.6 of yeast, or the protein was associated with a regulatory task (Supplementary Material 6) [17,41,42],

### Reconstruction of the extended stoichiometric model and general design considerations

Reconstruction involved the modification of 4 existing species (all metabolites), 13 species types (11 metabolites and 2 enzymes), and 53 existing reactions, as well as the removal of 3 reactions from Y7.6. 90 species types (68 enzymes, 18 metabolites and 4 pseudo-metabolites) representing 173 species (67 enzymes, 11 pseudo-metabolites and 95 metabolites), and 104 new enzymatic and transport reactions were introduced to the model. The new version has improved gene coverage of the existing model by 6%.

Reactions, which were reported to take place at the membrane but did not have a detailed mechanism explained, were represented to occur across the membrane for simplicity without compromising on the details of the iron metabolism. The enzymes associated with these reactions were localised to the membrane in order to highlight the nature of the transmembrane reaction. The pseudo-metabolites were introduced such that they did not interfere with the material, energy or the redox balance of the network. They were introduced in coupled reactions in order to avoid accumulation, and these reactions were unbounded so that the system was not constrained by the flux limitations through these reactions when the underdetermined system was optimised for a given objective.

Only 14 new dead-end metabolites could be detected indicating that the extension by incorporation of the iron metabolism did not disrupt the connectivity of the metabolic network. Although the Fe-S clusters matured in the nucleus are not recruited by the metabolic enzymes in the current reconstruction, this process was included nevertheless in order to allow future extensions in the model, at the cost of introducing phosphate and matured 4Fe-4S species as dead-end metabolites in the nucleus. Other dead-end metabolites were introduced through cyclic interconversions; PS (extracellular), NADPH, NADP, FMN, and FMNH_2_ (in vacuole), and FMNH_2_ (in the mitochondria). Some by-products of the iron metabolism are allowed to accumulate in the yeast cell as reported in the literature; bilirubin and ferrioxamineB (in vacuole), coprogen, ferrichrome, and enterobactin (in the cytoplasm). L-cysteine is transported into the mitochondrion in the existing model for the assembly of the sulfonylated scaffold for the Fe-S cluster formation. Although recent reports have suggested a putative cysteine synthase (Mcy1p) identified in the mitochondrial outer membrane [44], this mechanism has not yet been extensively investigated and therefore was excluded.

97% of the reactions through Y7.Fe were comparable to those of Y7.6 and only 850 *(ca.* 25%) reactions had non-zero fluxes when unit glucose uptake rate was employed as the single constraint for optimising growth. Of those 850 reactions, the flux through only 233 remained the same and the difference in fluxes through 266 of the remaining reactions was more than 25%. This indicated a substantial rewiring in more than 1/3^rd^ of the active reactions in the model under standard basic growth conditions upon introduction of the iron metabolism into the genome scale metabolic network model. The predicted growth flux was reduced by 7% in the extended model (Supplementary Material 3) indicating the cost of the iron metabolism on the existing network. The high metabolic burden was most likely to be associated with energy generation, which could not be captured by Y7.6, since this observation was in line with the growth predictions obtained by constraining the fluxes through the purine pathways in our preliminary analysis. Revisiting the same system, the flux predictions improved substantially employing Y7.Fe with only a ±20% difference between the experimentally measured and predicted growth fluxes, at the cost of lower predictive accuracy of the growth phenotype. Indeed, an analysis of those reactions, which displayed more than 25% change in their flux between Y7.6 and Y7.Fe, were associated with genes that were significantly enriched for the nucleoside phosphate metabolic process (p-value<10^−38^) along with other metabolic processes in line with our observations on the analysis involving the purine intermediates employed as flux constraints (Supplementary Material 3).

The predictive performance of the model was further investigated through gene essentiality. The extended model was observed to perform very similarly to the existing primary model with only marginal differences in measures evaluating predictive power of the model despite a sizeable improvement of 6% in gene coverage (Table 1). Major improvements and extensions in the metabolic network model structures were previously observed to have even more substantial effects on the predictive power as a trade-off [9]. The viability predictions for both models indicated a significant over-enrichment for the successful prediction of gene essentiality; 2.83-fold (p-value<10^−27^) and 3.04-fold (p-value<10^−30^) for Y7.Fe and Y7.6, respectively.

**Table 1.** Evaluation of the predictive power of the extended model

|  | **Y7.6** | **Y7.Fe** |
| --- | --- | --- |
| ***Number of genes*** | 908 | 963 |
| ***Number of TP*** | 677 | 709 |
| ***Number of TN*** | 83 | 84 |
| ***Number of FP*** | 73 | 90 |
| ***Number of FN*** | 76 | 80 |
| ***PPV (%)*** | 90 | 89 |
| ***NPV (%)*** | 52 | 48 |
| ***Sensitivity (%)*** | 90 | 90 |
| ***Specificity (%)*** | 53 | 48 |
| ***Predictive success (%)*** | 84 | 82 |

## Results and Discussion

### Iron-recruiting enzyme requirements of the metabolic network model

The stoichiometry by which the cofactors were introduced to the reactions in the metabolic network was observed to be closely related to the predictions on growth rate. We observed a tight control of how much the metabolic iron requirement of the network model was over the growth rate. The total iron requirement of the cell was calculated from the iron composition of a standard defined medium for yeast [45], and the recruitment of iron or complex iron entities as cofactors by the enzymes was simulated based on the constraint imposed by the iron transport flux. A fine-tuning for the sustainable iron recruitment indicated that the stoichiometric coefficients for these terms need to be in the order of magnitude of 10-14 without compromising substantially on the growth rate (Figure 2a). The metabolic requirement of total turnover for iron-recruiting enzymes was determined as 3.02 × 10^−11^ mM / (g biomass)^-1^ **h^−1^** based on this model. The maximum and the minimum theoretical requirement for iron-recruiting enzymes was determined by investigating the variability of these fluxes. This analysis demonstrated that the cell was employing the pathways where iron recruitment of the cognate enzymes would be required minimally, possibly to improve energy efficiency of the system. The total turnover was only 1.5% higher than that computed for the theoretical minimum usage. On the other hand, we observed that the fluxes through these pathways would be rewired such that the iron-recruiting enzyme turnover would be increased by 34% (Supplementary Material 3).

**Figure 2.**
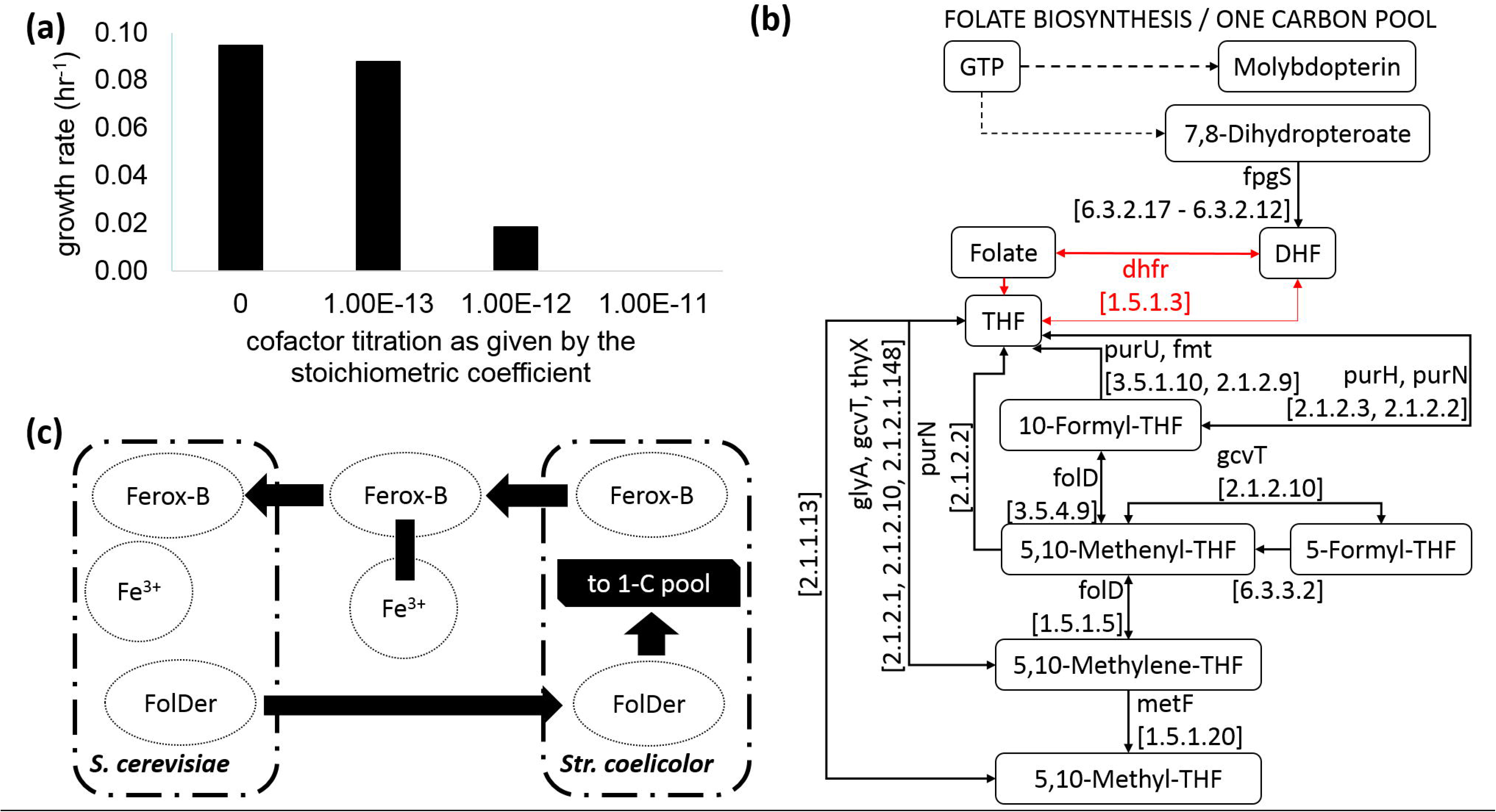
Indispensable role of iron for yeast and alternative routes for iron scavenging. The cellular requirement for iron is well documented for *S. cerevisiae* monocultures. This plot demonstrates how growth rate predictions of the metabolic network model is affected by the amount of iron recruited by the metabolic enzymes. The stoichiometric coefficient of iron ions and complex iron entities to be recruited by iron requiring enzymes without impairing growth substantially can be determined based on the constraints imposed by the experimentally permissible limits of iron uptake (a). A schematic representation of the folate biosynthesis and the one carbon folate pool in *Str. coelicolor* is given in (b). The metabolites are represented in rounded rectangles, arrows indicate the directionality of the reactions and enzymes for the reactions are indicated adjacent to the cognate arrows. Dashed lines indicate more than one reaction step lumped together. DHFR related routes are indicated in red. The EC numbers of the enzymes are provided in brackets. A possible scheme of sharing economies between *S. cerevisiae* and *Str. coelicolor* are given in (c). The directionality of the arrows indicate the suggested direction of material flow in a mutualistic coexistence event, not the directionality of the transporters. The blunt-end connection indicates binding of the siderophore to a ferric ion. Ferrox-B: ferrioxamine B, FolDer: folate and its derivatives, 1-C pool: one carbon folate interconversions pathway.

The stoichiometric coefficients for cofactors were determined as very low, as expected within its biological context. Although iron was not among the trace element cofactors, the stoichiometric coefficients were determined in the magnitude of 10^−14^. Working with such small numbers impose technical constraints for the analysis as well. The precision of the machine and the solver employed can both inflict problems at this stage. For the current problem, the precision of both the machine and Gurobi were two orders of magnitude lower than the stoichiometric coefficients, allowing us a robust platform to conduct this analysis independent of whether the MATLAB or the Python version of Cobra was utilised. However, for less abundant cofactors, this is an issue that needs to be kept under consideration.

### Benchmarking the model against environmental and genetic challenges

Having established the iron-recruiting enzyme requirement of the metabolic network, we extended the investigation to understand how the *in silico* system adapted to environmental challenges that were tailored specifically to exploit this model. The response of the network to different levels of extracellular iron and copper availability was investigated through flux distributions. For this purpose, the upper and lower bounds of the fluxes through several reactions were set to zero to mimic reports available in the literature [31,32], High-affinity iron system was activated by blocking the routes through low affinity Fe^2+^ uptake routes. The absence of extracellular xenosiderophores was also taken into consideration for the simulation of an *S. cerevisiae* monoculture. The predictions on flux distributions were in agreement with the expected physiological outcome on how the yeast metabolism responded to low or high abundance of copper [46] or iron [47],

We then investigated how the loss or reduction of functionality in *ARH1*, *ATM1* or *YFH1*; essential components of the ISC machinery, could be studied employing Y7.Fe. All three genes are essential although only *ARH1* could be identified as inviable employing the model. The essentiality of *YFH1*, the Fe-S cluster scaffold protein, or *ATM1*, the Fe-S apocluster transporter, is prompted by the fact that they are required to supplement Fe-S clusters to be recruited on essential non-enzymatic proteins, making the biogenesis and pre-processing of these Fe-S clusters an essential process for the cell, but not specifically for the metabolism. Therefore, essentiality of a protein could only be captured if that essential function were represented in the model. Despite their essentiality, loss of a single copy of these genes in the diploid BY4743 genetic background did not yield any significant growth defects (p-value < 0.01) with specific growth rate for all mutants remaining within = 0.43 ±0.01 h^−1^. These mutants did not display significant differences (p-value <0.01) with respect to their utilisation of glucose or ammonium as the carbon and nitrogen sources respectively, nor in their production of ethanol or glycerol. On average, the mutants were observed to consume more iron per unit optical density equivalent number of cells although this difference was not observed to be significant due to the high variance within replicates. Significant difference in the intracellular distribution of copper was observed between the mutants and the wild type. However, the intracellular measurements of haem, copper, reduced or total iron content of the mutants were not sufficient enough to reveal the metabolic or non-metabolic essential roles of their cognate genes (Supplementary Material 3).

Although both the seed model and the iron-incorporated model had a propensity to over-predict essentiality as discussed in earlier sections, only Y7.Fe had a slight but significant enrichment in over-representation of essential genes (1.06-fold, p-value<10^−3^). Consequently, the reactions with nonzero fluxes were more often catalysed by enzymes encoded by essential genes (2.32-fold over-enrichment, (p-value<10^−40^)) in Y7.Fe. In line with this and earlier observations, *ARH1* was widely associated with reactions having non-zero fluxes in the metabolic network. Since *ARH1* was an essential gene also for the model, the reorganisation of the fluxes in the network could only be investigated by introducing an *“in silico* reduction of function”. We employed flux as a proxy for the functional capacity of the enzyme catalysing that reaction and lowered the flux value by 50% to mimic a reduction in the function of the enzyme catalysing that reaction. Since *ARH1* was an essential gene for the model and the predicted growth rate was also reduced by 50% in response to reducing the flux through the reaction catalysed by *ARH1*, the expected outcome for the remaining non-zero-fluxes was to be reduced by 50%, or to remain unchanged. However, a considerable fraction of the fluxes (8%) were observed to be either rewired or had a change in magnitude other than by 50% reduction. 13% of the enzymes and transporters represented in Y7.Fe (963 in total) were affected by these changes. The rewiring of the fluxes was predominantly observed to involve the lipid metabolism (p-value<10^−24^). Furthermore, fluxes were observed to rewire away from the glycine and serine family amino acid metabolism (p-value<10^−6^) towards the neutral lipid metabolism (p-value<10^−4^). The enzymes associated with the reactions that displayed unexpected variations in the magnitude of their fluxes were significantly associated with the central carbon metabolism (p-valueclO^−6^), energy generation and respiration (p-value<10^−3^) and nucleotide metabolism (p-valueclO^−16^). Among the reactions with unexpectedly altered fluxes, 32% were orphan transport or exchange reactions. Many of these reactions were involved with the transport or exchange of lipid metabolism intermediates across organelles as well as of amino acids including glycine, L-alanine, L-leucine, L-threonine, serine, and valine, and of small inorganic molecules including ammonia, bicarbonate, carbon dioxide, hydronium ion, oxygen, phosphate, and water (Supplementary Material 3). The rewiring of the lipid metabolism and changes in the fluxes in the energy pathways in response to a perturbation induced to mimic the reduction of *ARH1* function in the cell was in line with earlier findings on its role in biological systems. Apart from acting as an essential component of the ISC machinery, *ARH1* is a protein homologous to the human adrenodoxin reductase [48], which was reported to function in the mitochondrial electron transfer chain that catalyses the conversion of cholesterol into pregnenolone. The retroviral expression of *ARH1* was shown to restore the LDL receptor function in cells from patients suffering from familial hypercholesterolemia [49], demonstrating the enzyme’s role in neutral lipid metabolism, as also captured by our analysis in the model yeast system.

### Yeast as the iron scavenger of the wilderness

The exhaustive search on iron associated enzymatic and transport routes highlighted the extensive siderophore-bound high-affinity iron transport and siderophore processing routes in *S. cerevisiae*. As the baker’s yeast has no endogenous siderophore production pathways available, this mechanism is dedicated solely to utilise xenosiderophores, which are likely to be present in yeast’s extracellular environment in the wilderness cohabitated and shared by other members of the community potentially producing and secreting these iron carriers. Although these routes were to be considered as inactive in all monoculture systems, and consequently in their model representations, the availability of routes for transporting xenosiderophore-bound iron had a broader implication with regards to yeast’s metabolic demand. The three questions that immediately arose were: (i) “Why has the baker’s yeast has conserved these alternative assisted routes of iron uptake to supplement its own direct uptake routes?”, (ii) “Which microbial species would potentially be involved in this activity?”, and (iii) “What determines the nature of yeast’s co-existence with another species in the community based on an iron dependent relationship?”.

Iron is an essential element for yeast and its metabolism has a higher demand for iron than any other cofactor, with 10% of the enzymes catalysing the metabolic reactions requiring iron in one form or another. A substantial proportion of these reactions lie at the core of the primary metabolism; in energy generation and de novo nucleotide biosynthetic pathways. This high demand bears the need for a constant supply of iron for growth and proliferation. We believe that it is precisely for this reason that yeast has evolved to have access to as many potential routes to allow iron influx as possible to ensure its survival in a mixed population. Yeast growth was shown to depend essentially on a constant influx of iron, and stoichiometric iron requirement and the availability of extracellular iron necessitated fine-tuning of this demand in the metabolic analysis of a monoculture (Figure 2a). This tight control of iron over growth addresses the first question as to why these potential assisted iron scavenging routes were conserved in yeast.

We addressed the next two questions together by looking for a fungal or prokaryotic species that would coexist with *S. cerevisiae* in nature with minimal incompatibilities. The KEGG Metabolic Pathways database, one of the largest resources available on metabolic networks, hosts data on the metabolic networks of approximately 110 fungal species and 4400 prokaryotic species [50]. Among this wide range of possibilities, we imposed constraints to narrow down our search space; (i) evidence for coexistence with *S. cerevisiae*, (ii) evidence for the endogenous production and export of at least one of the siderophores that *S. cerevisiae* can uptake: coprogen, ferrichrome, TAFC, enterobactin, or ferrioxamine B, (iii) availability of a curated genome-scale metabolic network model to allow an in silico study of the metabolism. We further limited the search space by looking for a mutualistic relationship between *S. cerevisiae* and the candidate species, where, in exchange for iron, the candidate organism would receive a metabolite from yeast, which it cannot synthesise or becomes limited by, thus creating a bottleneck for its metabolic network. *Str. coelicolor* was identified as such a candidate with evidence of coexistence with *S. cerevisiae* [51], producing and excreting ferrioxamine B [52,53], and has curated genome scale metabolic networks available [54,55],

An investigation of the metabolic network of *Str. coelicolor* indicated that it lacked the dihydrofolate reductase (DHFR) enzyme (Figure 2b). This could only be a misrepresentation problem since DHFR family members were previously reported to be difficult to annotate due to their similarity to pyrimidine dehydrogenase family members (Pfam01872), which were reported to be numerous in Actinomycetes like *Str. coelicolor* [56]. A protein BLAST [57] search for this highly conserved enzyme did not yield any recognised sequence on the *Str. coelicolor* genome, and the culture media commonly employed for the species is generally known to be supplemented with folate or folate derivatives apart from HMM, where the growth was always observed to be poor (Glyn Hobbs, personal communication), all hinting at the possibility that a functional DHFR might indeed be missing in *Str. coelicolor* and that other partially conserved functional domains in pyrimidine dehydrogenase family enzymes or in methylenetetrahydrofolate reductase were performing poorly to replace the role of DHFR. This could then potentially generate a potential bottleneck in the operation of the one carbon pool, which we propose to be a possible incentive to establish a mutualistic relationship between *Str. coelicolor* and *S. cerevisiae*. We hypothesised that in an environment where resources were limited, ferrioxamine B-bound iron could be scavenged by the yeast and the bacterium in exchange could benefit from the availability of folate and folate derivatives that yeast could synthesise. Folate transporters, which are available in both species, were previously reported to work bi-directionally [58], and be able to also recognise tetrahydrofolate [59], facilitating this sharing economy. Employing the genome scale metabolic network model for *Str. coelicolor* [55], we were able to remove the pseudo reaction to represent DHFR activity in the model and confirm the inability of the system to sustain growth in the absence of folate, and recuperate growth in the presence of folate uptake flux, which was set to remain within the maximal production rate by *S. cerevisiae*. Furthermore, we employed Y7.Fe to confirm inability to grow in the absence of iron, which was then recovered by the introduction of ferrioxamine B-bound iron to sanction the prospect of such a sharing economy (Figure 2c).

## Conclusion

The complex network structure comprised of the interactions among enzymes, metabolites and their regulators define transitions between metabolic states. Metabolic state shifts designate improved adaptation in response to environmental or genetic perturbations, which are desirable in some cases such as biotechnological applications, whereas adaptation would be most unwelcome in other situations such as switching to a diseased state such as to induce cancer. Studying the metabolic network at the genome scale allows us to achieve a better understanding of how the fluxes can get rewired in response to an internal or external input. This renders extensive and comprehensive network representation at the heart of analysis in addressing a variety of metabolic states and conditions. This paper presents a strategy to allow for a comprehensive extension of the genomescale stoichiometric network model of the yeast metabolism by incorporating a hugely underrepresented cellular system; non-metabolic ion cofactor involvement. We selected iron ion as our test case as it is involved in nearly 10% of all enzymatic steps in the metabolic network of yeast. Although the work is iron-specific, our analysis provided a new approach for handling co-enzymes and co-factors in stoichiometric models, which can be generalised for other such entities. We made use of pseudo metabolites, which were not produced or consumed by the cell, in modelling the system, since these intermediates functioning in a cyclic interconversion pathway allowed a more explicit description of the molecular mechanisms represented in the model. We identified the optimal turnover rate of iron cofactors in the metabolic network utilising this reconstruction, which is not possible to achieve experimentally employing the currently available analytical technologies. We also investigated the demand for iron cofactors in metabolism from an evolutionary perspective leading yeast to adopt alternative metabolic strategies to scavenge extracellular iron available for other co-existing species. Furthermore, an in-depth adjunct analysis of the *Str. coelicolor* metabolic network allowed us to identify potential bottlenecks involving DHFR.

It should also be noted that the final Y7.6Fe model is not intended be evaluated as an extended version of the stoichiometric model of the metabolic network due to the presence of pseudo metabolites, and enzyme cofactors introduced as substrates and products, which do not undergo a biochemical change. This model presents an unconventional way of extending the metabolic network models towards incorporating non-metabolic information, which could be valuable in incorporating non-metabolic cofactor involvement essential for constructing whole cell models of biological systems.

## Acknowledgments

Authors would like to thank Ayca Cankorur-Cetinkaya and Alexander P. S. Darlington for useful discussions on the copper-iron metabolisms in yeast, Glyn Hobbs for discussions on *Str. coelicolor* physiology, and Daniel J. H. Nightingale for discussions and guidance on the subcellular fractionation of yeast organelles.

## Funding Statement

Authors gratefully acknowledge the funding from the Leverhulme Trust (ECF-2016-681 to DD), EC 7th FP (BIOLEDGE Contract no: 289126 to SGO), BBSRC (BRIC2.2 to SGO).

## Supplementary material captions

**Supplementary Material 1** Extended model of the metabolic network of yeast (SBML level 2, version 4) (file format: .xml).

**Supplementary Material 2** Details on the extended model. The modifications carried out on the primary model, the new species types, and the new species including both genes and metabolites are provided in separate worksheets. The reactions that are introduced into the network are described and provided along with their gene associations accompanied by the logical rules for these genes whenever applicable, the reversibility rules, the literature/database evidence for the reaction, and the upper and lower bounds of each reaction in the last worksheet (file format: .xlsx).

**Supplementary Material 3** Worksheet on flux distributions and predictions on viability. The evaluation of the distribution of fluxes between the primary model and the existing model, the growth predictions on single gene deletions, double gene deletions, the distribution of fluxes for reduced flux through the reactions catalysed by Arh1p, and the flux variability analysis for determining the enzyme abundance upper- and lower-bounds are provided in separate tabs (file format: .xlsx).

**Supplementary Material 4** Table summarising the Fe-S cluster requirements of metabolic enzymes employed in the metabolic network model Yeast 7.6 (file format: .docx).

**Supplementary Material 5** Table summarising the iron family cofactor requirements of the enzymes and active transporters of the yeast metabolic network model (file format: .docx).

**Supplementary Material 6** Table summarising the enzyme-cofactor relationships excluded from this reconstruction and detailed reasons for exclusion (file format: .docx).

